# Reinforcement motor learning after cerebellar damage is related to state estimation

**DOI:** 10.1101/2023.08.17.553756

**Authors:** Christopher M. White, Evan C. Snow, Amanda S. Therrien

## Abstract

Recent work showed that individuals with cerebellar degeneration could leverage intact reinforcement learning (RL) to alter their movement. However, there was marked inter-individual variability in learning, and the factors underlying it were unclear. Cerebellum-dependent sensory prediction may contribute to RL in motor contexts by enhancing body state estimates, which are necessary to solve the credit-assignment problem. The objective of this study was to test the relationship between the predictive component of state estimation and RL in individuals with cerebellar degeneration. Individuals with cerebellar degeneration and neurotypical control participants completed two tasks: an RL task that required them to alter the angle of reaching movements and a state estimation task that tested the somatosensory perception of active and passive movement. The state estimation task permitted calculation of the active benefit shown by each participant, which is thought to reflect the cerebellum-dependent predictive component of state estimation. We found that the cerebellar and control groups showed similar magnitudes of learning with reinforcement and active benefit on average, but there was substantial variability across individuals. Using multiple regression, we assessed potential predictors of RL. Our analysis included active benefit, somatosensory acuity, clinical ataxia severity, movement variability, movement speed, and age. We found a significant relationship in which greater active benefit predicted better learning with reinforcement in the cerebellar, but not the control group. No other variables showed significant relationships with learning. Overall, our results support the hypothesis that the integrity of sensory prediction is a strong predictor of RL after cerebellar damage.

## INTRODUCTION

The cerebellum has long been known to play a critical role in controlling movement [1]. A dominant hypothesis is that the cerebellum houses internal models of body and environment dynamics, which it combines with information about motor commands to generate predictions of the sensory consequences of actions (i.e., the forward internal model) [2,3]. Sensory predictions serve as internal feedback signals that compensate for time-delayed peripheral afference, permitting rapid and well-coordinated movement. In the event of changes to the body or environment dynamics, a form of supervised motor learning, often called adaptation, is thought to recalibrate internal models (and sensory predictions) to keep them up to date [4,5]. A large body of literature supports this hypothesis, demonstrating how the motor symptoms of cerebellar damage correspond with disrupted internal model and sensory prediction calculation, as well as how cerebellar damage impairs adaptation (for work in humans, see [6–8] and [9–16], respectively). Impaired adaptation has been posited to underlie poor rehabilitation outcomes in this population [17–19].

Yet, motor learning is achieved through a combination of multiple neural mechanisms [20,21], and the effect of cerebellar damage on mechanisms other than adaption is less well understood. In recent years, the effect of cerebellar damage on reinforcement motor learning (RL) has received particular attention [22–24]. RL is a form of semi-supervised learning. While adaptation relies on continuous feedback about the direction and magnitude of movement errors, RL relies on periodic scalar measures of outcome [25,26]. At the simplest level, reinforcement feedback can take the form of binary signals indicating success or failure. RL requires individuals to explore possible task solutions to determine which one(s) yield(s) success. Over time, the probability of success is learned for each task solution, and this permits a biasing of motor selection to favor the solution(s) most likely to generate a positive outcome.

Therrien et al. (2016) [22] showed that individuals with cerebellar degeneration could use binary reinforcement to learn to rotate the angle of a reaching movement comparably to a group of age-matched control participants. The same individuals could not learn in adaptation conditions. In a later pilot study, Therrien et al. (2021) [27] showed that individuals with cerebellar degeneration could also use binary reinforcement to learn reduce elements of reaching ataxia. Together, these results suggested that interventions leveraging RL hold promise as a novel approach to rehabilitation training for cerebellar ataxia. However, there was variability in the degree of learning observed across individuals in both studies. While most individuals with cerebellar degeneration learned well in RL conditions, some individuals did not. Understanding the factors underlying inter-individual variability in task performance would improve the translation of RL interventions in this patient population, as it would permit screening of which individuals are most likely to benefit from treatment.

A mathematical model of the RL task by Therrien et al. (2016) [22] posited that efficient learning from binary reinforcement required accurate state estimation. In other words, to solve the credit assignment problem and map reinforcement signals to the correct movement, the motor system must know what action was produced. The model was agnostic to the cause of poor state estimation; however, the hypothesized role of the cerebellum in sensory prediction suggests that, in the case of cerebellar degeneration, variation in sensory prediction impairment may underlie differences in state estimation (and RL) across individuals. Indeed, others have proposed a similar idea using different models [23,28]. Sensory prediction is thought to contribute to state estimation [29,30]. The contribution is evidenced by an enhancement of somatosensory acuity during active (i.e., voluntary) movement when sensory predictions can be computed using motor command information, compared to passively induced movement when they cannot [31–33]. Cerebellar degeneration has been shown to disrupt this ‘active benefit’ [34–37]; however, Weeks et al. (2017a) [36] demonstrated that the degree of active benefit disruption can vary across individuals, with many showing comparable levels of active benefit to age-matched control participants. Indeed, the group level comparison of active benefit did not reach significance in their study. Variance in active benefit across individuals in the cerebellar group was not correlated with clinical impairment scores and was suggested to reflect different patterns and/or extent of cerebellar degeneration. It is conceivable, then, that the magnitude of active benefit shown by an individual with cerebellar degeneration could constitute a metric of the integrity of sensory prediction’s contribution to state estimation. A correlation between active benefit and RL across individuals with cerebellar degeneration would support the hypothesis that sensory prediction impairment predicts poor RL in this patient population.

Impaired predictive state estimation is a parsimonious predictor of inefficient motor RL after cerebellar damage, yet it is far from the only possibility. The severity of individuals’ ataxia, or their baseline movement variability, may impede the performance of the correct task solution regardless of whether an individual can discern what that is. Simply reducing movement velocity can additionally improve motor output in this population (it makes the movement dynamics easier to control and increases the time available to use delayed peripheral feedback) and, hence, permit better expression of learning [6,7,38]. Finally, learning with binary feedback has been shown to rely heavily on explicit strategy use in younger neurotypical populations [39,40]. Cerebellar damage has been implicated in the learning of explicit strategies (vs. their execution) [41,42], although there is evidence that participant age may pose a larger constraint on strategy computation [43]. Aging is also associated with reduced sensitivity to feedback in non-motor reward learning tasks [44–46]. Consequently, variation in RL across individuals could reflect age-related declines in cognitive capacity.

The purpose of this study was to assess the relationship between RL and active benefit in a sample of individuals with cerebellar degeneration. We first sought to replicate prior results by comparing individuals with cerebellar degeneration to a group of neurotypical control participants matched for age and biological sex as they performed an RL task that required them to alter the angle of reaching movements. Both groups also performed a state estimation task that tested the somatosensory perception of active and passive movement, permitting the calculation of active benefit. We then used multiple regression to test the significance of potential predictors of the magnitude of learning in the RL task. Our results showed that the cerebellar and control groups performed similarly in both tasks, on average, but there was substantial inter-individual variability. In the cerebellar group, the magnitude of learning in the RL task showed a significant relationship with the degree of active benefit in the state estimation task. This relationship was not significant in the control group. No other predictors showed significant relationships with learning in either group. We discuss the significance of these findings by considering how they contribute to an understanding of a cerebellar role in motor RL and the potential application of RL in rehabilitation training protocols for individuals with cerebellar damage.

## MATERIALS AND METHODS

### Participants

The study was approved by the Einstein Healthcare Network and Jefferson Health Institutional Review Boards, and all participants gave written informed consent before participation. A total of 12 people with cerebellar degeneration (CB; 5 female, 7 male, mean age 55.1 years, SD: 15.6 years) and 12 neurotypical individuals (Control; 5 female, 7 male, mean age 55.8 years, SD: 15.8 years) participated in the study. Further details about the CB group’s characteristics are shown in Table 1. CB participants were screened for and presented with no extra cerebellar or extrapyramidal signs on a neurological exam, normal passive peripheral sensation in the upper extremities, and no history of drug or alcohol abuse or other neurological conditions. One participant with cerebellar degeneration (CB05) presented with hyperreflexia in the lower limbs; however, reflexes in the upper limbs were within normal limits. Control participants were also screened for and presented with no history of drug or alcohol abuse or neurological condition. CB participants’ movement impairment severity was assessed using the International Cooperative Ataxia Rating Scale (ICARS) [47].

**Table 1.** Participant Demographics.

| Participant | Age<br>(years) | Sex<br>(M/F) | Handedness<br>(R/L) | Diagnosis | ICARS |  |
| --- | --- | --- | --- | --- | --- | --- |
|  |  |  |  |  | Total<br>(/100) | Limb Kinetic<br>(/52) |
| CB01 | 46 | F | R | ADCAIII | 41 | 17 |
| CB02 | 23 | M | R | SAOCA | 46 | 23 |
| CB03 | 75 | M | R | SCA14 | 50 | 23 |
| CB04 | 58 | F | L | SCA8 | 23 | 8 |
| CB05 | 59 | M | R | SAOCA | 52 | 28 |
| CB06 | 58 | F | R | ADCAIII | 24 | 7 |
| CB07 | 69 | F | R | SAOCA | 31 | 15 |
| CB08 | 37 | M | R | ADCAIII | 25 | 11 |
| CB09 | 55 | M | R | SAOCA | 42 | 23 |
| CB10 | 73 | M | R | Autoimmune | 45 | 16 |
| CB11 | 42 | M | R | SAOCA | 31 | 13 |
| CB12 | 66 | F | R | SAOCA | 55 | 20 |
| **CB Group** | **Mean: 55.1**<br>**SD: 15.6** | **M: 7**<br>**F: 5** | **R: 11**<br>**L: 1** |  | **Mean: 37.8**<br>**SD: 10.9** | **Mean: 16.6**<br>**SD: 6.3** |
| **Control Group** | **Mean: 55.8**<br>**SD: 15.8** | **M: 7**<br>**F: 5** | **R: 11**<br>**L: 1** |  |  |  |

CB: Participant with cerebellar degeneration. Control: Neurotypical. M: Male; F: Female. R: Self-reported right-hand dominant. L: Self-reported left hand dominant. ADCAIII: Autosomal dominant cerebellar ataxia type 3. SAOCA: Sporadic adult-onset cerebellar ataxia. SCA: Spinocerebellar ataxia types 8 and 14. ICARS: International Cooperative Ataxia Rating Scale. The ICARS Total reflects the total score out of a possible 100. The ICARS Limb Kinetic reflects the limb coordination sub-score out of a possible 52. Higher scores on the ICARS indicate more severe ataxia.

### Apparatus

The experimental tasks were performed using the KINARM Exoskeleton robot (Kinarm, Kingston, Ontario, Canada). The Kinarm restricted the motion of the arms to the horizontal plane. Hands, forearms, and upper arms were supported in troughs appropriately sized to each participant’s arms, and the linkage lengths of the robot were adjusted to match the upper limb segment lengths for each participant to allow smooth motion at the elbow and shoulder. Vision of the arms was obstructed by a horizontal mirror, through which participants could be shown targets and cursors in a veridical horizontal plane via an LCD monitor (60 Hz refresh rate). Movement of the arm was recorded at 1 kHz.

### Procedure

Participants completed two tasks with their self-reported dominant arm: a reinforcement motor learning task and a somatosensory state estimation task. The tasks were performed in separate sessions, and the order was fixed so that participants completed the reinforcement motor learning task first.

#### Reinforcement Motor Learning (RL) Task

Figure 1a shows a schematic of the task. This task was adapted from Therrien et al. 2016 [22] and required participants to learn to rotate the angle of 10 cm, target-directed reaching movements in a counterclockwise direction by 15 degrees. Prior work showed that individuals with cerebellar degeneration exhibit a clockwise bias when reaching to a target at 90-degrees (i.e., the end target used here) with a Kinarm Exoskeleton robot [48]. Therefore, we only tested the counterclockwise version of the task to ensure that participants would not perform well simply because of their bias and would be required to learn a new reach angle to achieve task success. The reach angle was defined as the angle between two vectors: one from the center of the home position to the center of the target and another from the center of the home position to the index fingertip location when it crossed a radial distance of 10 cm from the center of the home position (i.e., the reach endpoint). Before performing the task, participants were given time to practice making the target-directed reaching movements with and without visual feedback of their arm to familiarize themselves with the Kinarm and task environment. In the experimental task, all participants performed a 70-trial baseline phase with no rotation. Visual feedback of the hand position was provided for the first 10 trials of the baseline phase and then removed for the remaining 60 trials. Only data from the 60 baseline trials without visual feedback was used for analysis of baseline performance. Following the baseline phase, participants completed a 180-trial perturbation phase in which they learned to rotate the reach angle. Participants were given a short break (1-3 min) every 60 trials throughout the task to avoid the effects of fatigue.

**Figure 1.**
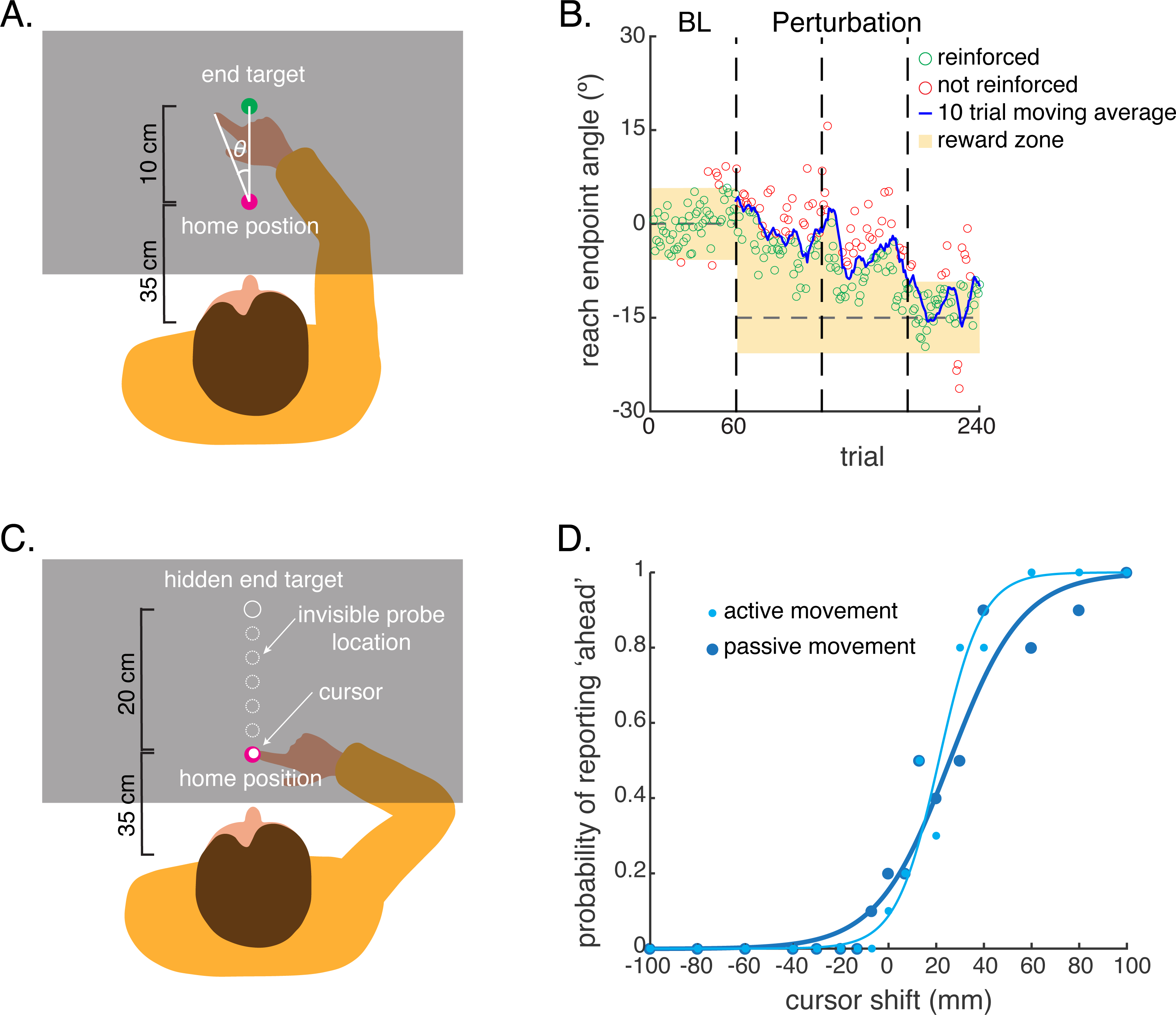
A schematic of the reinforcement motor learning (RL) and somatosensory state estimation tasks. Both tasks were performed with the dominant arm. **(a)** The RL task had participants make 10 cm reaching movements from a home position to a single end target. The task drove participants to learn to rotate the endpoint angle of their reaches. The reach angle was defined as the angle between one vector connecting the home position to the end target and another connecting the home position to the index fingertip position when it crossed a radial distance of 10 cm from the home position (i.e., the reach endpoint). Throughout the task, the arm moved below a screen and was hidden from view. Participants received binary feedback about the outcome of their reaches. Reaches that successfully crossed a reward zone were reinforced with the target turning green and an auditory tone. Reaches that did not cross the reward zone resulted in the target turning red and no auditory tone. **(b)** Data from an example individual with cerebellar degeneration showing the trial and feedback structure of the RL task. After a baseline phase in which the reward zone corresponded to the visually displayed target, closed-loop reinforcement feedback was used to drive participants to rotate their reach angle. A running average of the reach angle from the previous 10 reaches was computed. Reaches with an angle that fell between the running average and the rotated target were reinforced. Reach angles that fell outside this range were not reinforced. **(c)** In the somatosensory state estimation task, participants actively moved or were passively moved along a 20 cm reach trajectory. A visual cursor showed the index fingertip position for a brief period at the beginning of each trial before being extinguished. On each trial, the hand was stopped at 1 of 5 possible probe locations. A visual cursor reappeared for a 600 ms display period, but its position was shifted relative to the true position of the index finger. After 600 ms, the cursor disappeared, and participants completed the reach. Participants then reported whether they perceived the cursor as ‘ahead’ of or ‘behind’ the index fingertip during the display period in a 2-alternative forced choice paradigm. **(d)** Data from the same example participant shown in panel B. The probability of reporting ‘ahead’ was determined for each of the 17 cursor shift values tested. The data from each movement condition were then fit with separate psychometric functions, from which measures of somatosensory acuity and bias were derived.

A trial began with participants’ index fingertips in a home position (a pink colored circle with 1 cm radius) located 35 cm in front of the chest and aligned to the sagittal midline of the body. A cursor (a white circle with 0.5 cm radius) was projected over the index fingertip. Once the arm was held in that position for 500 ms, the cursor was extinguished, and the end target appeared on the screen (a light-blue colored circle with 1 cm radius). The center of the end target was positioned 10 cm distal to the home position, directly in front of it. Participants were instructed to begin their reach when they saw the target appear and to reach so that the index fingertip passed through the center of the end target. The trial ended when the index fingertip exceeded a radial distance of 10 cm from the center of the home position; the index fingertip position at this point was used to calculate the reach angle in real time. After a post-trial delay of 700 ms, the target was extinguished, and the participant could move their hand toward their chest. Once the index fingertip was within a 6 cm radius of the home position, the fingertip cursor reappeared to help participants reposition the arm for the next trial. To encourage participants to complete their movement within 200-600 ms after leaving the home position, the text “Too Fast” or “Too Slow” was displayed on the screen in the post-trial delay period following reaches that fell outside the desired speed range.

Throughout the task, participants received binary (i.e., success or failure) feedback about the outcome of their reaches. Reaches with an angle that fell within a reward zone were reinforced with the end target changing color from light blue to green and an auditory tone (Figure 1b). Reaches with an angle that fell outside the reward zone resulted in the end target changing color from light blue to red and no auditory tone. Participants were not informed about the component of their reaches mapped to the feedback signals. That is, they were not explicitly instructed to alter their reach angle. Rather, they were instructed to use the binary feedback provided to discern the correct reaching movement throughout the task. Participants were, thus, required to explore different reaching movements to discover the one that was reinforced.

During the baseline phase, the reward zone corresponded to the visually presented target – i.e., the index finger had to pass through the visually displayed target. During the perturbation phase, a closed-loop reinforcement schedule (Figure 1b) was used to drive participants to rotate the endpoint of the reach in a counterclockwise direction [22]. With closed-loop reinforcement, the reward zone was determined by calculating a moving average of the reach angles from the previous 10 trials. The reward zone (i.e., the yellow-shaded region in Figure 1b) corresponded to reach angles between the moving average and the rotated target. As participants learned to reach at angles that fell in the reward zone, their 10-trial moving average shifted accordingly, narrowing the reward zone and progressively pushing participants to rotate their reach angle to the new target position.

#### Somatosensory State Estimation Task

This task was adapted from Weeks et al. 2017b [37]. It required participants to report the location of a visual cursor relative to the perceived position of the unseen index fingertip in a 2-alternative forced choice paradigm. The reports were made after voluntary reaches (i.e., the active movement condition) or after reaches were passively induced by the robot (i.e., the passive movement condition). Reaches were performed with the dominant arm along the y-axis of the workspace to make them comparable to the movements performed in the RL task. Somatosensory perception was assayed at the reach endpoint because the RL task required participants to learn to bias the position of the reach endpoint.

Figure 1c shows a schematic of the task. All trials began with the robot passively moving the arm to place the index fingertip in a home position (a pink colored circle with 1 cm radius). A visual cursor (a white circle with 0.5 cm radius) then displayed the index fingertip position for 2 seconds before being extinguished along with the home position. Showing the true index fingertip position at the beginning of each trial served to realign somatosensory perception and prevent sensory drift. In the passive movement condition, the robot moved the hand to one of five possible probe locations along a 20 cm reach path. In the active movement condition, an auditory “go” signal cued participants to reach voluntarily until they encountered a haptic wall that stopped the reach at one of the five possible probe locations. The five probe locations prevented participants from habituating to a single repeated arm movement. Once at the probe location, the visual cursor reappeared for a 600 ms display period, in which its position was shifted relative to the index fingertip. Seventeen shift values were tested: ±100, ±80, ±60, ±40, ±30, ±20, ±13, ±7, 0 mm. Following the display period, the cursor was extinguished. Participants then completed the remainder of the 20 cm reach, either passively or actively, depending on the movement condition. In the active movement condition, the reach endpoint was signaled by a second haptic wall. At the reach endpoint, participants verbally reported whether they perceived the cursor had been “ahead” of or “behind” their index fingertip during the display period. The experimenter recorded the participants’ reports, and then the robot passively moved the arm back to the home position to begin the next trial.

In the active movement condition, a 5 cm wide force channel was used to constrain reach trajectories to the desired region of the workspace. The width of the channel ensured that participants could not rely entirely on the haptic walls to guide their reach and perform the task correctly. This prevented them from using the channel wall as an additional source of sensory information in the task. Participants were encouraged to keep their average movement speed between 7 – 27 cm/s in the active movement condition. The text “Too Fast” or “Too Slow” was displayed on the screen at the end of trials when movement speeds fell outside this range. In the passive movement condition, participants were moved at 17 cm/s corresponding to the slowest permittable movement speed in the reinforcement motor learning task.

The probe location and shift magnitude were randomized across trials in both movement conditions. Using the method of constant stimuli, participants completed 10 repetitions of each shift magnitude, corresponding to 2 repetitions of each shift magnitude per probe location and a total of 170 trials per movement condition. The order of passive and active movement conditions was counterbalanced across participants.

### Analysis

All kinematic analysis was performed using custom-written scripts in Matlab (Mathworks), and statistical analysis was performed using SPSS v29 (IBM).

In the RL task, reach angles were computed from the raw position data and expressed relative to the target location so that angles in the clockwise direction were positive and angles in the counterclockwise direction were negative. To compare RL task performance between groups, the dependent variables of mean reach angle, reach angle standard deviation, and reach peak velocity were assessed using mixed analyses of variance (ANOVA). Between-subjects comparisons were made across a factor of group (cerebellar degeneration, CB; neurotypical, Control), and within-subjects comparisons were made over two experimental phases: the final 20 trials of the baseline phase and the final 20 trials of the perturbation phase. Total learning – i.e., the difference between the end of the perturbation and baseline phases – was also compared across groups using an independent samples t-test. Within each group, the total learning was compared against zero using one-sample t-tests. Finally, the change in peak velocity between the baseline and the end of the perturbation phases was compared across groups using an independent samples t-test and compared to zero within groups using one-sample tests. Change values below zero indicated that participants increased their speed from baseline to the end of the perturbation phase, whereas positive values indicated that they slowed down.

For the somatosensory state estimation task, the proportion of trials in which a participant reported the cursor to be “ahead” of the index fingertip was determined for each shift value. The data were then fit with a psychometric (logistic) function [49]. Separate functions were fit for each subject and movement condition. Three dependent variables were derived from the fitted psychometric functions: the just noticeable difference (JND), the active benefit, and the bias. The JND quantified the slope of the psychometric function and served as a measure of somatosensory acuity. The JND was calculated for each subject and movement condition as the difference in shift values (i.e., the x-axis in Figure 1d) corresponding to the 75% and 25% probabilities (i.e., the y-axis in Figure 1d) on the psychometric function. A smaller JND indicated better somatosensory acuity. We calculated active benefit as the difference in JND between the active and passive movement conditions. Values below zero indicated active benefit (i.e., the active JND was smaller than the passive JND). The bias quantified the midpoint of the psychometric function and served as a measure of the point of subjective equality (i.e., the shift value at which participants perceived the cursor to be aligned with the index fingertip position). The bias was calculated as the shift value corresponding to a 50% probability on the psychometric function. The statistical analyses of JND and bias served to compare task performance across groups and used mixed ANOVA with a between-subjects factor of group (CB, Control) and a within-subjects factor of movement condition (passive, active). Pearson correlation was used to assess the relationship between JND and bias across movement conditions. The active benefit was compared across groups using an independent samples t-test and compared against zero within groups using one-sample t-tests.

Within the active movement condition, additional analyses were performed to compare movement speed and trajectory variability across groups. Specifically, independent samples t-tests were used to compare the reach peak velocity and the lateral variability of reach movements (i.e., the standard deviation of movement along the x-axis) between the Control and CB groups. We also assessed the bivariate correlations (Pearson r) between these measures and the active benefit within each group.

Finally, we used multiple linear regression analysis to assess predictors of reinforcement motor learning in the CB and Control groups. We constructed a total of 5 regression models. Total learning in the RL task was the dependent variable in all models. Two regression models were used to test predictors of RL in the CB group. The first model included the following independent variables: the limb kinetic sub-score of the ICARS (i.e., the severity of CB participants’ limb ataxia), the baseline reach angle variability in the RL task, the change in peak reach velocity in the RL task, participants’ age, and the active benefit shown in the somatosensory state estimation task. The second model controlled for the possibility that active benefit values were driven by one movement condition in the state estimation task. This model included the limb kinetic sub-score of the ICARS, the baseline reach angle variability in the RL task, the change in peak reach velocity in the RL task, participant’s age, and the JND values from the active and passive movement conditions of the somatosensory state estimation task. The third and fourth regression models tested predictors of RL in the Control group. These models followed the same construction as the first two models, with the exception that they did not include the ICARS score as an independent variable. The fifth regression model was constructed as part of a *post hoc* analysis and tested the significance of the interaction between group and active benefit as a predictor of RL. This final model included the independent variables of group, active benefit, and the group-by-active-benefit interaction term.

All data were tested for normality using Shapiro-Wilk tests. Homogeneity of variance was assessed using Mauchly’s Test of Sphericity. To ensure that data met the assumptions for multiple regression analysis, we used the Durbin-Watson test for independence of observations, assessed the Tolerance/VIF values for multicollinearity, and the normal P-P plot for approximately normally distributed residuals. For all statistical analyses, the alpha value was set to 0.05.

## RESULTS

Figure 2A shows the time series of reach angles for the cerebellar degeneration (CB) and neurotypical (Control) groups in the reinforcement learning (RL) task. Analysis of variance (ANOVA) was used to compare the mean reach angle between groups and across two task phases (the final 20 trials baseline and the final 20 trials of the perturbation phases). This analysis showed a significant main effect of task phase (*F*(1,22) = 22.696, *p* <.001) in which the mean reach angle at the end of the perturbation phase was significantly lower at baseline. Although the CB group learned slightly less, there was no main effect of group (*F*(1,22) = .684, *p* = .417) or group-by-phase interaction (*F*(1,22) = .824, *p* = .374). This result was corroborated by analysis of the total learning (Figure 2B), which was not significantly different between groups (*t*(22) = -.908, *p* = .374), and both groups showed total learning that was significantly greater than zero (CB: *t*(11) = 2.645, *p* = .023; Control: *t*(11) = 4.142, *p* = .002).

**Figure 2.**
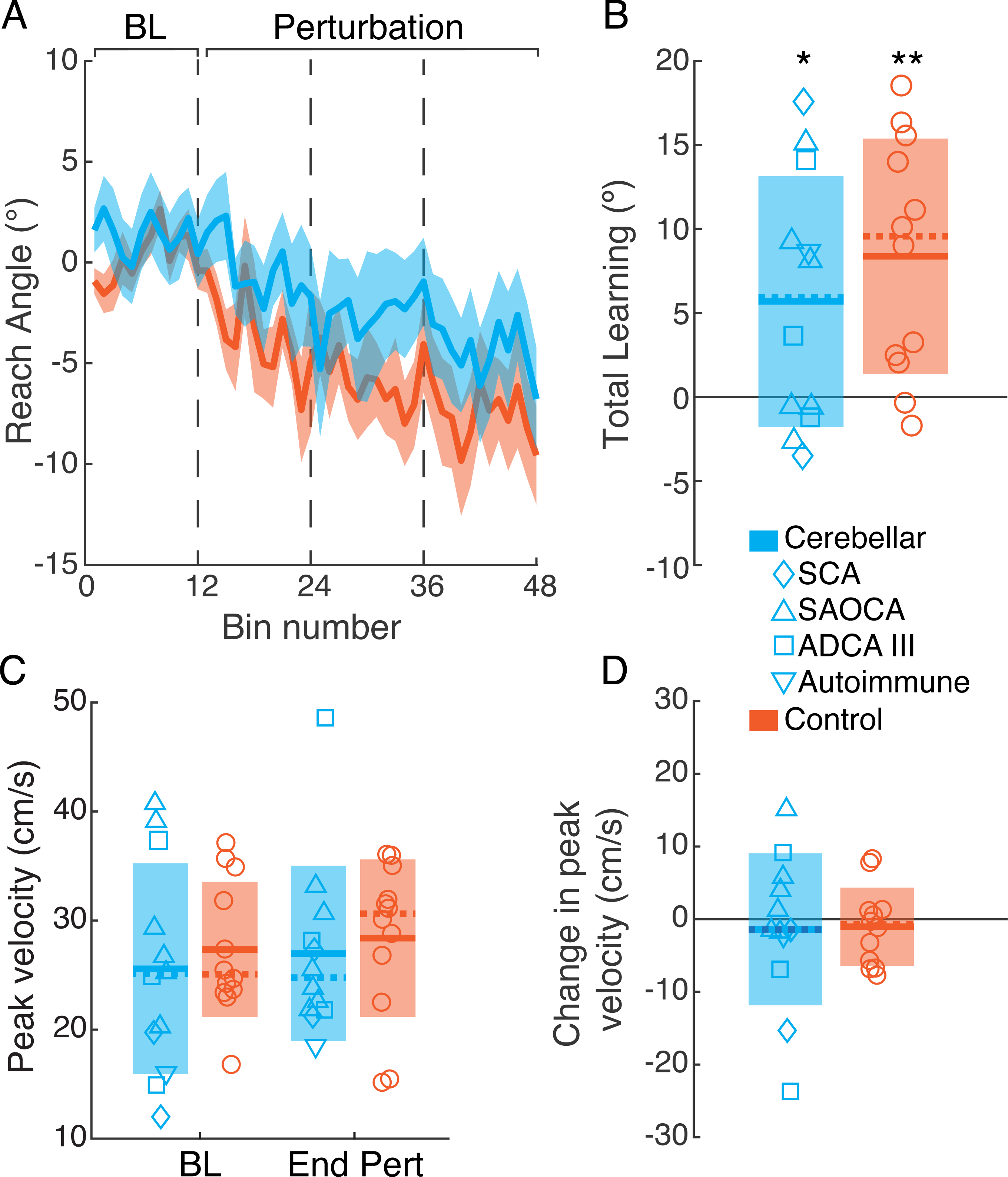
Behavior in the reinforcement motor learning task. (a) The time series of reach angles shown by the cerebellar and control groups. Solid lines represent the group mean, and the shaded regions depict the standard deviation. Each bin comprises the mean of 5 trials. (b) The total learning (i.e., the difference in mean reach angle between the end of the baseline and perturbation phases) shown by each group. (c) The mean reach peak velocity shown by each group at the end of the baseline (BL) and perturbation (End Pert) phases. (d) The within-subject change in reach peak velocity between the end of the baseline and perturbation phases. (b-d) Markers represent individual participants. Solid lines represent the group mean. Dashed lines represent the group median. The shaded regions depict the standard deviation. SCA: Spinocerebellar ataxia. SAOCA: Sporadic adult-onset ataxia. ADCA III: Autosomal dominant cerebellar ataxia type 3. Autoimmune: Autoimmune mediated cerebellar degeneration. * *p* <.05 *\*\* p* <.01.

ANOVA testing differences in reach angle variability (i.e., standard deviation) between groups and across task phases showed no significant main effects of group (*F*(1,22) = .452, *p* =.566) or task phase (*F*(1,22) = 3.814, *p* = .064), or group-by-phase interaction (*F*(1,22) = .460, *p* = .505). Similarly, ANOVA comparing reach velocity across the groups and task phases showed no significant main effects or interaction (Figure 2C; group: *F*(1,22) = .340, *p* = .566; task phase: *F*(1,22) = .516; group-by-phase: *F*(1,22) = .012, *p* = .914). To quantify changes in reach velocity within subjects, we computed the difference in reach peak velocity between the baseline and end of perturbation phases (Figure 2D). The CB group showed larger between-subject variance than the Control group, with most CB participants reducing their speed over the course of the RL task. However, the difference between groups was not significant (*t*(22) = -.110, *p* = .914), and neither group distribution was significantly different from zero (CB: *t*(11) = -.465, *p* = .651; Control: *t*(11) = -.667, *p* = .519).

Figure 3 shows the performance of both groups in the somatosensory state estimation task. ANOVA comparing the just noticeable difference (JND) values across groups and movement conditions showed a significant main effect of movement condition (*F*(1,22) = 7.559, *p* = .012; Figure 3A). The effect was driven by smaller JND values, indicating improved somatosensory acuity, in the active compared to the passive movement condition. There was no main effect of group (*F*(1,22) = 2.262, *p* = .147) or group-by-movement-condition interaction (*F*(1,22) = .017, *p* = .898), which indicated that both groups showed comparable performance. The JND values in the active movement condition showed a significant positive correlation with JND values in the passive movement condition within both groups indicating that participants performed consistently in both conditions (CB: *r* = 0.783, *p* =.003; Control: *r* = 0.922, *p* <.001). That is, individuals with greater acuity in the active movement condition also showed greater acuity in the passive movement condition and vice versa.

**Figure 3.**
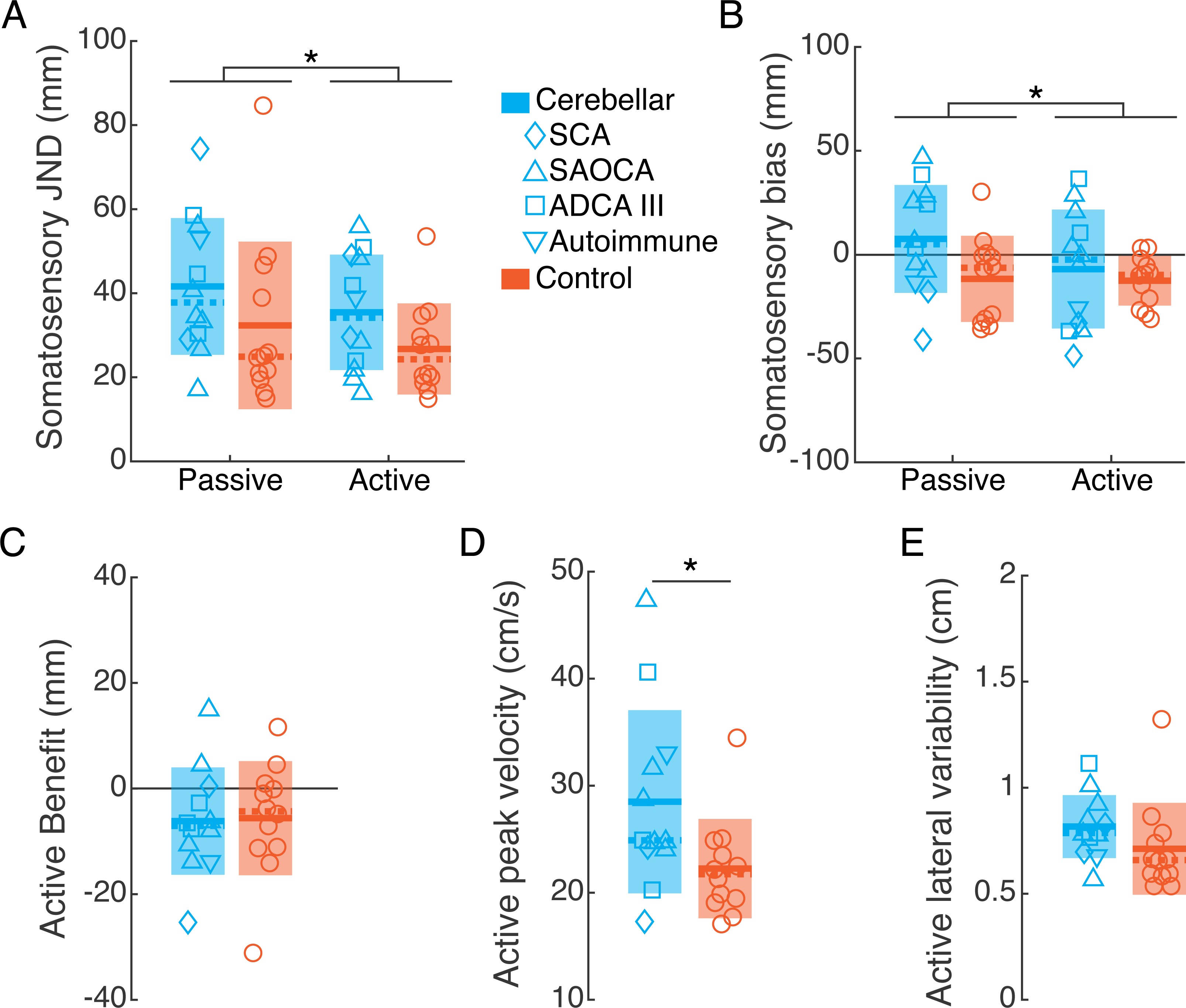
Behavior in the somatosensory state estimation task. (a-b) The just noticeable difference (JND, a) and the bias (b) shown by the cerebellar and control groups in the passive and active movement conditions. (c) The active benefit (i.e., the within-subject difference in JND between the active and passive movement conditions) shown by each group. (d) The mean peak velocity of reaching movements in the active movement condition shown by each group. € The standard deviation of position along the x-axis of reaching movements in the active movement condition shown by each group. (a-e) Markers represent individual participants. Solid lines represent the group mean. Dashed lines represent the group median. The shaded regions depict the standard deviation. SCA: Spinocerebellar ataxia. SAOCA: Sporadic adult-onset ataxia. ADCA III: Autosomal dominant cerebellar ataxia type 3. Autoimmune: Autoimmune mediated cerebellar degeneration. * *p* <.05

The results of the JND analysis were supported by the analysis of the active benefit (i.e., the difference in JND between the active and passive movement conditions) shown by each participant (Figure 3C). Means comparisons showed no significant difference in active benefit between groups (*t*(22) = .129, *p* = .898), and neither group distribution was significantly different from zero (CB: *t*(11) = 2.101, *p* = .059; Control: *t*(11) = 1.799, *p* = .100).

ANOVA comparing the somatosensory bias across groups and movement condition showed a main effect of movement condition, in which there was an overall negative bias in the active movement condition compared to approximately zero bias in the passive movement condition (Figure 3B; *F*(1,22) = 5.236, *p* = .032). There was no significant main effect of group (*F*(1,22) = 2.094, *p* = .162) or group-by-movement-condition interaction (*F*(1,22) = 4.056, *p* = .056), suggesting that both groups showed a comparable pattern of bias across the two movement conditions. Within the cerebellar group, there was a significant positive correlation between bias in the active and passive movement conditions (*r* = 0.891, *p* <.001); however, the Control group did not show a consistent bias across conditions (*r* = 0.458, *p* = .210).

To quantify any differences in movement kinematics between groups in the active movement condition, we analyzed the peak velocity and the lateral trajectory variability (i.e., the standard deviation of position in the x-axis) of each reaching movement. Means comparisons showed that the CB group moved with greater velocity than the Control group on average (*t*(22) = 2.222, *p* = .037). However, the peak movement velocity did not show any relationship with the active benefit in either group (CB: *r* = .189, *p* = .557; Control: *r* = .033, *p* = .918). Despite the width of the force channel being somewhat large (5 cm), both groups showed a comparable degree of lateral movement (*t*(22) = 1.383, *p* = .180). Lateral movement variability also showed no relationship with active benefit (CB: *r* = .032, *p* =.920; Control: *r* =.188, *p* = .559).

The primary objective of this study was to assess the relationship between a measure of internal model integrity and reinforcement learning in the CB group. In particular, we aimed to assess the significance of this relationship in the presence of other potential predictors of learning. To this end, multiple linear regression was used to test if the following independent variables significantly predicted total learning in the RL task: the active benefit shown in the somatosensory state estimation task, the severity of CB participants’ limb ataxia, the baseline variability shown in the RL task, the change in reach velocity in the RL task, and CB participants’ age. The overall regression was statistically significant (R2 =.807, *F*(5,6) = 5.014, *p* = .037). Of all the independent variables included in the model, only the active benefit significantly predicted total learning in the RL task (*β* = -.783, *p* = .007; Figure 4). The complete results of this regression analysis are shown in Table 2. To control for the possibility that the effect of active benefit may have been driven by one movement condition in the somatosensory state estimation task, we conducted a second regression analysis that included the active and passive JND values as independent variables in place of active benefit. Limb ataxia severity, baseline reach variability, the change in reach velocity, and participant’s age were included as control predictors. In this analysis the overall regression did not reach significance (R2 = .719, *F*(5,6)=3.077, *p* = .102), suggesting that none of the independent variables sufficiently captured the variance in total learning.

**Figure 4.**
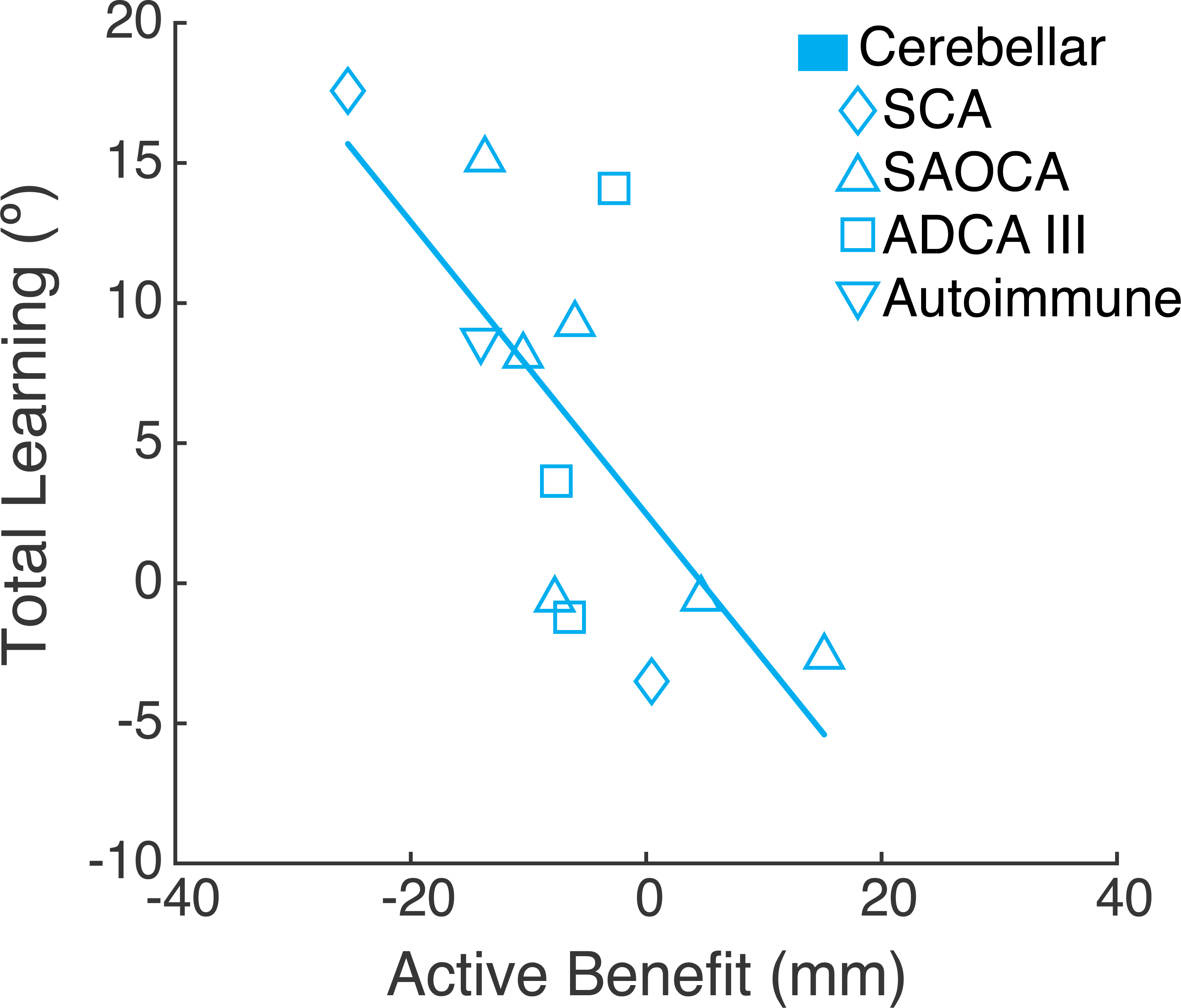
The relationship between the magnitude of active benefit in the somatosensory state estimation task and the total learning in the reinforcement learning task in the cerebellar group. The regression line depicts the bivariate regression for illustrative purposes. See Table 2 for the complete results of the regression analysis performed. SCA: Spinocerebellar ataxia. SAOCA: Sporadic adult-onset ataxia. ADCA III: Autosomal dominant cerebellar ataxia type 3. Autoimmune: Autoimmune mediated cerebellar degeneration.

**Table 2.** Results of multiple linear regression analysis to assess predictors of reinforcement learning in participants with cerebellar degeneration.

| Predictor | $\beta$ | Std. Error | $t$ | $p$ |
| --- | --- | --- | --- | --- |
| ICARS limb kinetic score | .357 | .216 | 1.650 | .150 |
| Baseline reach variability in the RL Task | -1.290 | .999 | -1.291 | .244 |
| Change in reach peak velocity in the RL task | -.228 | .131 | -1.736 | .133 |
| Participant age | -.251 | .127 | -1.969 | .097 |
| Active benefit | -.783 | .197 | -3.968 | .007 ** |
| Constant | 15.056 |  |  |  |
| R<sup>2</sup> | .807 |  |  |  |
| $F$ | 5.014 | | | |
| SEE | 4.435 |  |  |  |
| n | 12 |  |  |  |

\*\* $p < .01$ . ICARS: International Cooperative Ataxia Rating Scale. RL: Reinforcement learning.

Using multiple regression, we additionally examined predictors of total learning in the RL task within the Control group. We constructed two regression models that followed the same design as those used to analyze the CB group, with the exception that clinical ataxia severity was not included as a predictor. Neither regression model reached significance (model 3: R2 = .122, *F*(4,7) = .242, *p* .906; model 4: R2 = .188, *F* (5,6) = .277, *p* = .910). Together, these results indicate that none of the independent variables tested sufficiently explained the variance in total learning across Control participants. When taken with the CB group results, they further suggest an interaction effect of group and active benefit on total learning in the RL task, wherein active benefit is a significant predictor of learning only in the CB group. However, post hoc regression analysis with group, active benefit, and the group-by-active-benefit interaction term as independent variables and total learning as the dependent variable did not reach the threshold of significance (R2 = .316, *F*(3,20) = 3.076, *p* = .051), reflecting that the study was not powered to test this interaction effect.

## DISCUSSION

In this study, we examined reinforcement motor learning (RL) and somatosensory state estimation in individuals with cerebellar degeneration (CB participants) as well as a group of neurotypical control participants (Control participants) matched to CB participants for age and biological sex. We found that the CB group learned slightly less than the control group in the present task, but the difference was not statistically significant. Within the CB group, there was inter-individual variability, with many participants learning well and others learning less well. Overall, this replicates our prior results [22]. The somatosensory state estimation task was modeled after the task used by Weeks et al. (2017b) [37] and was designed to permit calculation of the active benefit shown by each individual. Active benefit describes the improvement in somatosensory acuity that is typically seen when estimating active (i.e., voluntary) compared to passively induced movement of the body [31–33] and is thought to be subserved by cerebellum-dependent sensory prediction [35–37]. Here, CB and control groups showed comparable active benefit in the somatosensory state estimation task, which runs counter to some previous work [35], but replicates the results of Weeks et al. (2017a) [36]. Importantly, we found variance in the degree of active benefit across individuals. We found that active benefit was a significant predictor of RL in the CB group. The individual JND values from the active and passive movement conditions of the state estimation task did not show significant relationships with learning, which demonstrated that it is not simply somatosensory state estimation, but differences in sensory prediction that best predicts inter-individual variance in RL after cerebellar damage. Other independent variables assessed – namely, ataxia severity, movement variability, the change in movement speed, and participant age – also showed no significant relationship with learning. Overall, our results suggest that RL, while intact after cerebellar damage, is related to predictive state estimation in this population.

Yet, there remain additional factors that could explain total learning variance across CB participants in our RL task. Studies of younger neurotypical individuals have shown that motor learning with binary reinforcement is supported in large part by the use of explicit cognitive strategies [39,40]. Although, it should be noted that RL has also been shown to support an implicit process in samples with similar demographics [50–52], suggesting that learning with binary feedback likely reflects a mix of active ingredients. Cerebellar damage has been shown to disrupt the computation of explicit strategies in comparable motor learning tasks [41,42], but this disruption is difficult to disentangle from general cognitive declines associated with the advanced age of these study samples [43]. While we did not find a significant relationship between participant age and RL here, we did not quantitatively assess strategy use. It would be interesting for future work to directly test the relationship between explicit strategy computation and motor learning with binary reinforcement in individuals with cerebellar damage.

Variability in RL across CB participants has also been suggested to stem from a cerebellar role in the fundamental processing of reward information [53]. Indeed, there is evidence from studies of animal models that the cerebellum encodes information about appetitive and social rewards [54–57]. Studies in humans have also shown that cerebellar damage can disrupt learning from monetary rewards in non-motor tasks [58–60]. However, important differences exist between the cerebellar encoding of reward and the canonical reward encoding in midbrain dopaminergic neurons [54,61]. Furthermore, the relationship between elements of appetitive and monetary reward processing (e.g., reward utility, expected value computation, etc.) and learning with arbitrary binary feedback signals (e.g., an auditory tone) is unclear. Do individuals need to find binary feedback signals as fundamentally rewarding as monetary gain in order to effectively learn from them? Further studies are needed to answer this question.

Within the Control group, none of the regression analyses performed showed significant results. This finding suggested that none of the independent variables tested were significant predictors of RL in neurotypical participants. While the lack of a significant relationship between active benefit and RL in Control participants limits any inference of causality in the relationship between sensory prediction and RL, it does not rule out the possibility. The present study was designed to test the hypothesis that the predictive component of state estimation (as measured by active benefit) is a strong predictor of variance in RL across individuals with cerebellar degeneration. Our results showed this relationship. Although Control participants can also show variance in the predictive component of state estimation, this may not predict inter-individual variability in learning to the same degree. The intact nervous system may be able to compensate for variance in predictive state estimation in ways that the damaged nervous system cannot (e.g., [62]). Our finding of a trending interaction in which the predictive relationship between active benefit and RL was present only for the cerebellar group and not the control group supports this idea.

The active and passive movement conditions of the somatosensory state estimation task differed in more ways than just the contribution of sensory prediction. The haptic wall, which served as a stop signal for each probe location in the active movement condition, may have provided additional sensory input that aided performance. Prior literature has shown that individuals with cerebellar degeneration are able to use haptic feedback to compensate for their impairment [48]. Participants also had to monitor their movement speed in the active movement condition, which increased attentional demands. Despite these differences, we found significant positive correlations in task performance across the two conditions, particularly in the CB group.

Participants showed consistent behavior, such that individuals with high somatosensory acuity (i.e., a lower JND) in the passive movement condition also showed high acuity in the active movement condition, and vice versa. Furthermore, CB participants showed a consistent pattern of bias across the two movement conditions. One would not expect these correlations in performance if the differences outlined above had an outsized influence on the active movement condition.

The present results raise an interesting point regarding the potential utility of RL in designing novel motor rehabilitation therapies for CB participants. Our results suggest that the factor predicting inefficient RL after cerebellar damage – impaired sensory prediction – is the same factor thought to underly impaired adaptation in this population. Impaired adaptation is posited to be an important contributor to the variable rehabilitation outcomes seen in individuals with cerebellar damage [17–19]. Thus, it may seem disadvantageous to study interventions leveraging RL. However, Therrien et al. (2016) [22] showed that many CB participants could learn with binary reinforcement but could not learn in adaptation conditions (evidenced by the respective presence and absence of after-effects). This suggests that there may be a gradient in dependence on sensory prediction across learning mechanisms [63,64]. The implication for CB participants is that there may be a critical period in which internal model function has degraded enough to preclude adaptation but is sufficiently spared to permit relatively efficient RL. A pilot study by Therrien et al. (2021) [27] showed that an RL intervention could improve the irregular trajectories that characterize cerebellar reaching ataxia. Although studies with larger sample sizes and additional statistical modeling are needed, it is possible that a test of active benefit could one day serve as a metric of which CB participants might benefit from an RL intervention.

## CONCLUSION

This study provides evidence that effective reinforcement motor learning (RL) after cerebellar damage is related to the reliability of sensory prediction in state estimation. We compared a sample of individuals with cerebellar degeneration (CB participants) and a sample of age- and sex-matched neurotypical controls (control participants) as they performed an RL task and a task that tested somatosensory estimates of limb state after active and passive movement. The somatosensory state estimation task permitted assessment of active benefit – the degree of improvement in somatosensory acuity with active compared to passive movement – in each participant. The active benefit is thought to reflect the contribution of sensory prediction to state estimation. Overall, the two groups performed similarly in both tasks. However, there was marked inter-individual variance, which aligned with prior literature. Within the cerebellar group, inter-individual variance in RL was best predicted by active benefit, suggesting a relationship between cerebellar-dependent sensory prediction and RL.

## DECLARATIONS

### Ethical Approval

The study was approved by the Einstein Healthcare Network and Jefferson Health Institutional Review Boards. All participants gave written informed consent to participate and to publish the data collected.

### Authors’ Contributions

AST conceived and designed the research, analyzed data, prepared figures, interpreted results, and wrote the manuscript. AST and ECS programmed the experimental tasks. CW and ECS performed the experiments and assisted with data analysis.

### Availability of Data and Materials

All experimental data and analysis scripts are available for download here: https://osf.io/s86bp/?view_only=c85f779a6f734825be1950dd54145fe0

